# Label-free, fast, 2-photon volume imaging of the structural organization of peripheral neurons and glia in the enteric ganglia

**DOI:** 10.1101/2022.06.15.496266

**Authors:** Doriane Hazart, Brigitte Delhomme, Martin Oheim, Clément Ricard

## Abstract

The enteric nervous system (ENS), sometimes considered as a ‘second brain’ due to its large autonomy from central circuits is made of interconnected plexuses organized in a mesh-like network lining the gastrointestinal tract. Originally described as a leading actor of the regulation of digestion, bowel advance and intestinal secretion, its implication in various neuropathologies has recently been demonstrated. However, with few exceptions, its morphology and functions have been studied on thin sections of the intestinal wall or in dissected explants. Due to its intricate morphology, precious information on its three-dimensional (3-D) architecture and connectivity is often lost. In this context, we have developed a fast, label-free 3-D imaging method of the ENS, based on intrinsic signals of the tissue. We adapted a fast tissue-clearing protocol based on a high refractive-index aqueous solution, and then characterized the autofluorescence signals arising from the various cellular and sub-cellular components of the ENS. Immunofluorescence and spectral recordings complete this characterization. Finally, we demonstrate the fast acquisition of detailed 3-D image stacks of unlabeled mouse intestine, across the whole intestinal wall and including both the myenteric and submucosal enteric nervous plexuses using a new spinning-disk two-photon microscope. The combination of fast clearing (less than 15min for 73 % transparency), autofluorescence imaging and rapid volumetric imaging (less than a minute for the acquisition of s z-stack of 100 planes (150*150 µm) at 300-nm spatial resolution) paves the way for new applications in fundamental and clinical research.

## INTRODUCTION

The enteric nervous system (ENS) is often being described as a “2^nd^ brain” (Avetisyan et al., 2015; Schneider et al., 2019). Comprised of more than 100 million neurons organized in interconnected ganglia, the ENS is responsible of many intestinal functions including bowel mobility, mucosal secretion, modulation of endocrine secretion as well as blood-flow regulation. In addition, albeit being a quasi-autonomous part of the nervous system, the ENS has extensive connections with both the sympathetic and parasympathetic systems and it is engaged in a bi-directional signaling with the CNS (Costa et al., 2000; Avetisyan et al., 2015; Schneider et al., 2019).

Located in the small-intestine wall, the ENS is made up of two interconnected types of *plexuses*. The myenteric *plexus* (also known as *plexus* of Auerbach’s) is located in between the circular and longitudinal *muscularis*, two layers of smooth muscle cells that are responsible of the motility of the bowel. Neurons of the myenteric plexus mainly regulate the motor function of the gut. The submucosal plexus (aka. plexus of Meissner’s) is located in vicinity of the gut *lumen*. It mainly regulates secretory functions (Kulkarni et al., 2018; Schneider et al., 2019), (**Figure 1**). Both *plexuses* are composed of different neuronal subtypes as well as enteric glial cells that act on specific functions, interact with one another and they target specific effectors (Boesmans et al., 2013; Schneider et al., 2019).

**Figure 1:**
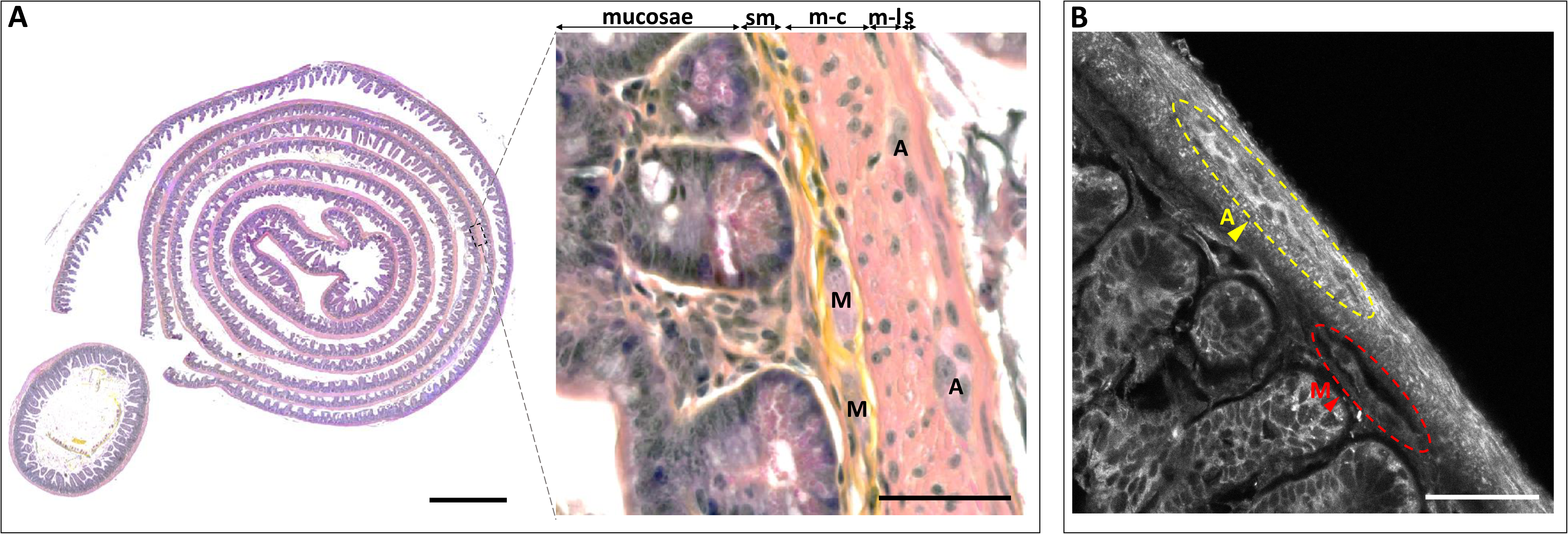
Histological observation of the ENS. (**A**) Hematoxylin/eosin/saffron stained intestinal Swiss roll (left) and transversal slice (left-inset). Magnification of a part of the intestinal wall observed on the Swiss roll and highlighting the different layers and components of the ENS (right). sm: submucosa, m-c: muscularis circular, m-l: muscularis longitudinalis, s: serosa, A: Auerbach’s plexus, M: Meissner’s plexus. Scale-bars: 500 µm (left), 50 µm (right). **(B)** Autofluorescence imaging from an intestinal Swiss roll under a 405nm excitation. A: Auerbach’s plexus, M: Meissner’s plexus. Scale-bar: 50 µm.

Abnormalities of the ENS have been linked to many pathologies of the digestive tract. One prominent example is Hirschsprung’s disease where regions of the large intestine (colon) lack ganglia. Without these nerve cells stimulating gut muscles to move contents through the colon, the contents can back up and cause constipation. More surprisingly, ENS abnormalities can be linked to central neuropathologies. For example, the analysis of the ENS by routine colonoscopy biopsies was shown to be a pre-mortem diagnostic tool of Parkinson’s disease, and studying ENS alterations provides insight into the progression of motor and non-motor symptoms, too (Lebouvier et al., 2010b; Ohlsson and Englund, 2019; Schneider et al., 2019). In addition, ENS dysfunction has been linked to other pathologies, including autism (Israelyan and Margolis, 2018; Heiss and Olofsson, 2019), thus underpinning the interest of a detailed morphological characterization of alterations of the cellular architecture of Meissner’s and Auerbach’s *plexuses* with modern biological imaging techniques.

Traditionally, anatomical or pathological studies of the ENS have relied on tissue slicing and conventional histological staining with hematoxylin and eosin (H&E), followed by imaging single thin sections with bright-field microscopy. Detailed volumetric observations, on the other hand, are rare and they mainly comprise very limited tissue volumes. This is, in part, because full 3-D reconstructions rely on first sequentially slicing the tissue bloc into thin sections, followed by their immunolabelling and sequential observation by confocal microscopy. Although routine in many fields of cell biology (where the sample volumes are much smaller), this immuno-confocal approach is extremely time-consuming for intestinal samples that measure between 0.3 mm (for a mouse colon) or mm for human biopsies: immunofluorescence protocols already require many hours and even days for the incubation of such thick samples. Also, by its scanning approach and point-wise image reconstruction, confocal laser-scanning microscopy (CLSM) is inherently slow for reconstructing large volumes. Furthermore, as the entire sample volume is illuminated but fluorescence only collected only from a single spot, confocal imaging rapidly photobleaches the entire volume during the repeated scans of a sample at different depths.

Two-photon (2P) excitation fluorescence microscopy (Denk et al., 1990) has become the gold standard for imaging thick tissue. Reduced in-plane photobleaching and enhanced depth penetration (Denk and Svoboda, 1997) combined with the efficient collection of scattered fluorescence (Oheim et al., 2001) are among its advantages for 3-D microscopy. Hundreds of fluorophores have action cross-sections that make them amenable to 2P imaging (Ricard et al., 2018) and tissue autofluorescence (AF) can be used to generate intrinsic contrast and provide context information (see, e.g., (Zipfel et al., 2003)). 2P microscopy and AF detection have been used for imaging various parts of the gut, including the stomach (Rogart et al., 2008), the small intestine (Orzekowsky-Schroeder et al., 2011; Ricard et al., 2012) and the colon (Aggarwal et al., 2013). However, these studies used scanning 2PEF microscopy that - like CLSM - is slow, often requiring 6 to 8 s/frame of intestine (see, e.g., (Ricard et al., 2012)) and thus several minutes for imaging a single block of intestinal sample.

Multi-spot 2P-imaging techniques parallelize either fluorescence excitation and/or pixel readout to overcome the low frame rate of single-spot scanning schemes (Bewersdorf et al., 2006). The most common of these are 2-P multi-spot scanning systems (Kim et al., 1999; Nielsen et al., 2001); and 2-photon spinning-disks microscopes (Egner and Hell, 2000; Straub et al., 2000; Kobayashi et al., 2002). Both have in common that they associate a planar fluorescence excitation through non-linear confinement with a “one-shot” readout on a fast camera detector. By design, these systems compromise on optical sectioning and image contrast, as cross talk between neighboring pinholes and scattered fluorescence becomes an issue, even at moderate imaging depths (see, e.g., (Shimozawa et al., 2013) for discussion).

Light-sheet fluorescence microscopy (LSFM) on the other hand has revolutionized the field of fast 3-D imaging. Here, a thin sheet of light orthogonal to the microscope axis is generated either by their cylindrical optics and/or scanning the beam in a plane. This planar excitation optically slices the sample. Fluorescence is collected perpendicular to the illuminated plane and imaged onto a camera. This configuration results combines improved contrast and axial sectioning with short acquisition and it reduces phototoxicity because only the imaged plane is illuminated at a given time (Huisken et al., 2004). However, even in its non-linear variant using pulsed scanned beams for planar 2-P excitation (Truong et al., 2011; Mahou et al., 2014; Zong et al., 2015), light-sheet microscopes suffer from a reduced *z*-resolution compared to spot-scanning systems. Also, LSFMs only work well on transparent samples, they require sample access from several sides, and often call for tedious embedding and mounting procedures.

In response to these challenges, we introduced a single-objective lens microscope scheme that retains the inline geometry of the classical microscope, boosts its speed beyond that of a light-sheet microscope and provides a significantly better z-resolution (Rakotoson et al., 2019). The optical sectioning, high acquisition speed and low photo damage are achieved through spatially and temporally changing 2PEF excitation to create a ‘virtual’ light sheet by aid a unique, single spinning-disk bearing both microlenses and pinholes. A high-numerical aperture (NA) low-magnification objective lens is used to image the excitation spots into the focal plane and to collect 2PEF. Our instrument combines the speed, field-of-view and ease of a 2-P light-sheet microscope with the resolution and image contrast of a scanning 2-P microscope. We previously demonstrated simultaneous dual-colour 3-D imaging of brain organoids with 300-nm lateral and 2.75-µm axial resolution over large fields of view at an imaging depth of 200 μm and with high temporal resolution (Rakotoson et al., 2019). Under realistic (low mW) excitation powers and signal levels typical for AF imaging or the detection of faint biological labels, a 900 × 900 pixel dual-color image was captured in 480 ms. These characteristics allow applications in 3-D morphological imaging of healthy and pathological thick biological samples.

One major drawback common to all parallelized 2-P imaging systems is that they work best with transparent samples or in superficial tissue layers. The reason is that a major advantage of single-spot 2-P imaging is sacrificed by using an imaging detector rather than a ‘bucket’ detector just collecting photons - the possibility of making use scattered fluorescence. Therefore, recent efforts in improving and speeding up volume imaging have typically gone along with improvements in tissue clearing (reviewed in (Richardson and Lichtman, 2015; Ueda et al., 2020; Tian et al., 2021; Weiss et al., 2021).

In the present work, we apply recent fast tissue clearing and volumetric imaging strategies for imaging the mouse small intestine. We demonstrate a label-free, highly contrasted 3-D imaging of enteric ganglia and an autofluorescence (AF-) -based identification of neurons and glial cells of the ENS. Our paper is organized as follows: (*i*), we first characterize a fast clearing approach of intestine sample compatible with the conservation of AF signals. (*ii*), characterizing AF spectra and intensity at a cellular and sub cellular scale, we identify neurones and enteric glial cells in both Auerbach’s and Meissner’s plexuses; (*iii*), we validate this AF-based identification of by confocal immunofluorescence. Finally, (*iv*), using our 2PEF OASIS microscope we demonstrate detailed volumetric imaging of the ENS throughout the entire depth of the intestinal wall. The here proposed workflow associating fast clearing and 3-D imaging will be a potential game-changer for biomedical applications, e.g., during diagnostic imaging.

## RESULTS

### Conventional histological observation is ill-suited for studying the 3-D organization and connectivity of the enteric nervous system

In clinical neurogastroenterology, as well as in fundamental research, the enteric nervous system (ENS) is mainly being studied using histological techniques that involve consecutive slicing, staining and observation steps. Briefly, in these approaches, the intestine is first fixed, then dehydrated, embedded in paraffin, sliced on a microtome to typically <10-µm thin sections, which are collected on a slide, rehydrated, stained with H&E and they are finally observed one-by-one on a light microscope. This approach is tedious, time-consuming and it risks– particularly in the case of a sparse 3-D structure like the ENS – to be little representative of the full biological complexity of the sample, as only a very small proportion can be observed. For example, on a typical transverse slice of the mouse intestine, it is very rare to find both Auerbach’s and Meissner’s plexuses, and the connections in between them are lost (**Figure 1A** - *left panel, inset*).

Alternatively, an approach dubbed “Swiss roll” largely increases the probability of finding both plexuses (Bialkowska et al., 2016). Here, a long stretch of intestine is opened longitudinally before rolling it around a tooth pick with the mucosa facing the internal part of the roll. The thus obtained, much longer sample is then processed as before for conventional histological fixation, embedding, slicing and staining. The use of the Swiss roll technique allows an about 10-fold gain in terms of the surface of the intestinal wall observable on a single slice (**Figure 1A –** *left*). The probability of finding both nervous plexuses within the same field of view is increased correspondingly (**Figure 1A – zoom**), however, despite its advantages, it is neither possible to observe the ENS in its full 3-D complexity, nor to study neuronal networks without a tedious serial slicing and reconstruction protocol.

Notwithstanding these limitations, we stained the such obtained Swiss-roll ENS section with classical hematoxylin/eosin/saffron (**Figure 1A – right panel**), an approach that can only be applied to thin slices to provide a benchmark image against which we will have to hold our new method. On the other hand, in preliminary experiments, we observed tissular autofluorescence (AF) – a signal arising from intestinal structures without any staining - and that appeared to produce sufficient contrast for identifying both plexuses (**Figure 1B**). We set out to study if this AF detection, with appropriate 3-D imaging techniques, is amenable to a full volumetric reconstruction of intact (= non-sliced) intestinal samples.

### A fast clearing protocol compatible with autofluorescence imaging

In intact intestinal samples the imaging depth is limited by absorption (mainly through pigments) and, in practice much more, by light scattering through the tissue. Tissue clearing techniques enhancing the transparency of the sample and homogenize its refractive index (RI). Together with advances in 3-D microscopy, tissue transparisation techniques have permitted detailed volumetric reconstructions of large tissue volumes. However, when it comes to reconstructing large 3-D volumes at and high 3-D spatial resolution, time-intense imaging and rendering sessions add to the already lengthy clearing and staining protocols (see, e.g., (Rakotoson et al., 2019)). To significantly facilitate the workflow of volumetric tissue imaging, we recently developed a ultra-fast, universal, non-toxic clearing protocol, based on a high-refractive index (RI) aqueous solution that was successfully tested on various organs (patent pending). However, the AF-based 3-D imaging of the ENS requires clearing to have a minimal attenuating effect on this already faint fluorescence signal. We therefore compared two protocols, one involving a depigmentation step of 45 min, the other one not. Using intact intestinal samples of 3 mm by 3 mm by10 mm, we compared the efficacy of the clearing on depigmented and non-depigmented tissue. In either case, the whole intestine became transparent and the depigmentation had little if any detectable effect on the final transparency (**Figure 2A**). We substantiated this observation by the quantification of the evolution with time of Michelson’s contrast (visibility) of a black and white stripe pattern placed underneath the intestinal sample in depigmented *vs*. non-depigmented conditions (**Figure 2B**). In either condition, a plateau was reached in less than 15 minutes and the final transparency of the samples was undistinguishable (71% *vs*. 73%; **Figure 2C - right panel**). The only parameter that was significantly different was the half-time for clearing (2 min 36 s depigmented *vs*. 5 min 18 s non-depigmented; **Figure 2C – left panel**). However, this tiny advantage seems negligible in view of the extra 45-min required for the depigmentation process. In addition, AF images of Auerbach’s plexus acquired upon 405-nm excitation revealed a stronger signal and higher contrast without depigmentation (**Figure 2D**). We conclude that depigmentation of the intestine is dispensable and that clearing of the whole intestinal wall can be performed in less than 15 min without destroying the autofluorescence signal.

**Figure 2:**
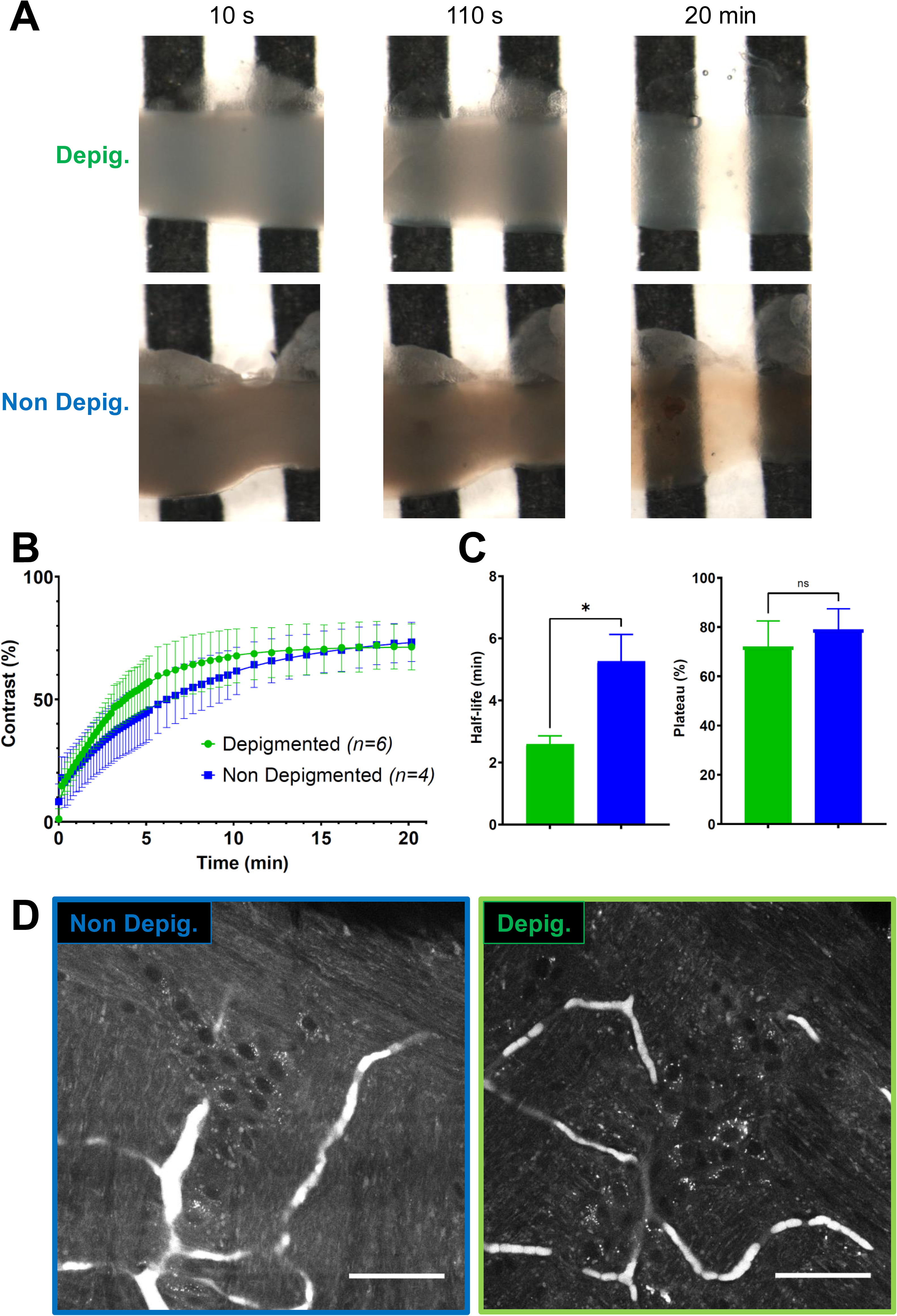
Fast clearing of the small intestine. (**A**) Longitudinal sections of ileum observed at various delays after the addition the clearing reagent (10 s, 110 s, 20 min) on a black and white pattern with (top) or without (bottom) a depigmentation step. (**B**) Quantification of the enhancement of the contrast in depigmented (green) and non-depigmented (blue) conditions. **(C)** Half-life (left) and efficiency of the clearing protocol (right) in depigmented (green) and non-depigmented (blue) conditions. (**D**) Autofluorescence imaging at the level of the Auerbach’s plexus under a 405nm excitation in non-depigmented (left) and depigmented (right) conditions. Scale-bar: 50 µm. Error-bars: standard deviation. *: p<0.05, Mann-Whitney test; ns: non-significant.

### Autofluorescence imaging allows a detailed morphological characterization of the ENS

We next confirmed that the morphologically identified AF structures were indeed neural cells. Intact intestinal samples were immunolabeled against HuC/D, a pan-neuronal marker (Mazzoni et al., 2021) and against S100β, an enteric glia marker (Grundmann et al., 2019). At the level of the muscularis longitudinalis, we detected only AF (**Figure 3 – first line**). However, within Auerbach’s plexus (**Figure 3 – second row**), neuronal cell bodies were located in the structures identified as ganglia on the AF images. Neurons exhibited strongly autofluorescent granules in their cell bodies. We could observe enteric glia in both ganglia as well as in the connective tracks in among them. Such connections were also clearly identifiable on the AF images despite their lower-intensity signal. Likewise, we could reliably observe neurons in the ganglia identified as Meissner’s plexus (**Figure 3 – third row**), albeit at lower density. Neurones in Meissner’s plexus exhibited less AF granules than those found in Auerbach’s plexus, and enteric glia were not present at this level. Finally, crypts exhibited solely AF signal (**Figure 3 – bottom row**).

**Figure 3:**
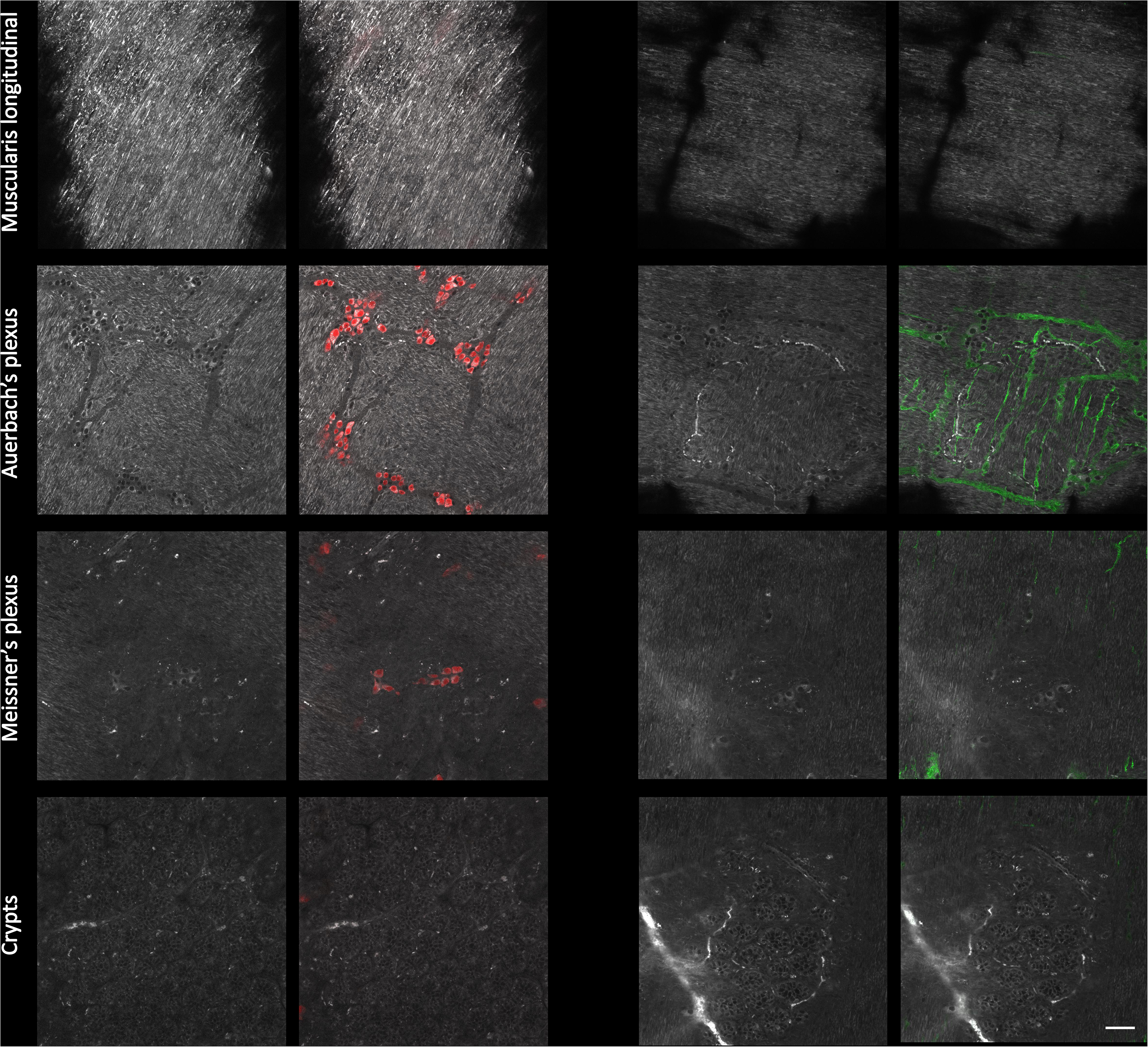
Immunohistological characterization of the autofluorescent structures of the ENS. Single autofluorescent images (grayscale) taken at different depth in the intestinal wall (muscularis longitudinalis, Auerbach’s plexus, Meissner’s plexus, crypts) and corresponding immunolabeled images. Red: neurons cell bodies (Hu C/D); green: enteric glia (S100β). Scale-bar: 50 µm.

### Spectral characterization of ENS autofluorescence

Tissue AF is generally a complex multi-component signal characterized by fairly large fluorescence excitation and emission spectra. We next wanted to know if ENS structures exhibited a specific AF signature that could be used to further pinpoint specific cellular and subcellular components, and we acquired AF emission spectra upon 405-nm excitation.

Within Auerbach’s plexuse*s*, regions of interest (ROIs) were selected in 4 different structures: (*i*), neuronal cell bodies that exhibited a strong granular AF (red ROI); (*ii*), cytoplasmic regions of enteric glia (green); (*iii*), neuronal nuclei (blue) and, (*iv*), ROIs within the muscularis (yellow) (**Figure 4A – left panel**). No significant differences were observed between the AF emission spectra for these 4 regions (**Figure 4B – left**). However, they can be discriminated based on the AF intensity that is strongest in neurons cell, followed by enteric glia, muscularis and nuclei that show the lowest AF. Moreover, neuronal cells bodies can also clearly be defined by the presence of numerous highly AF granules in their cytoplasm. Nuclei were characterized by a weak uniform signal (**Figure 4C – left**).

**Figure 4:**
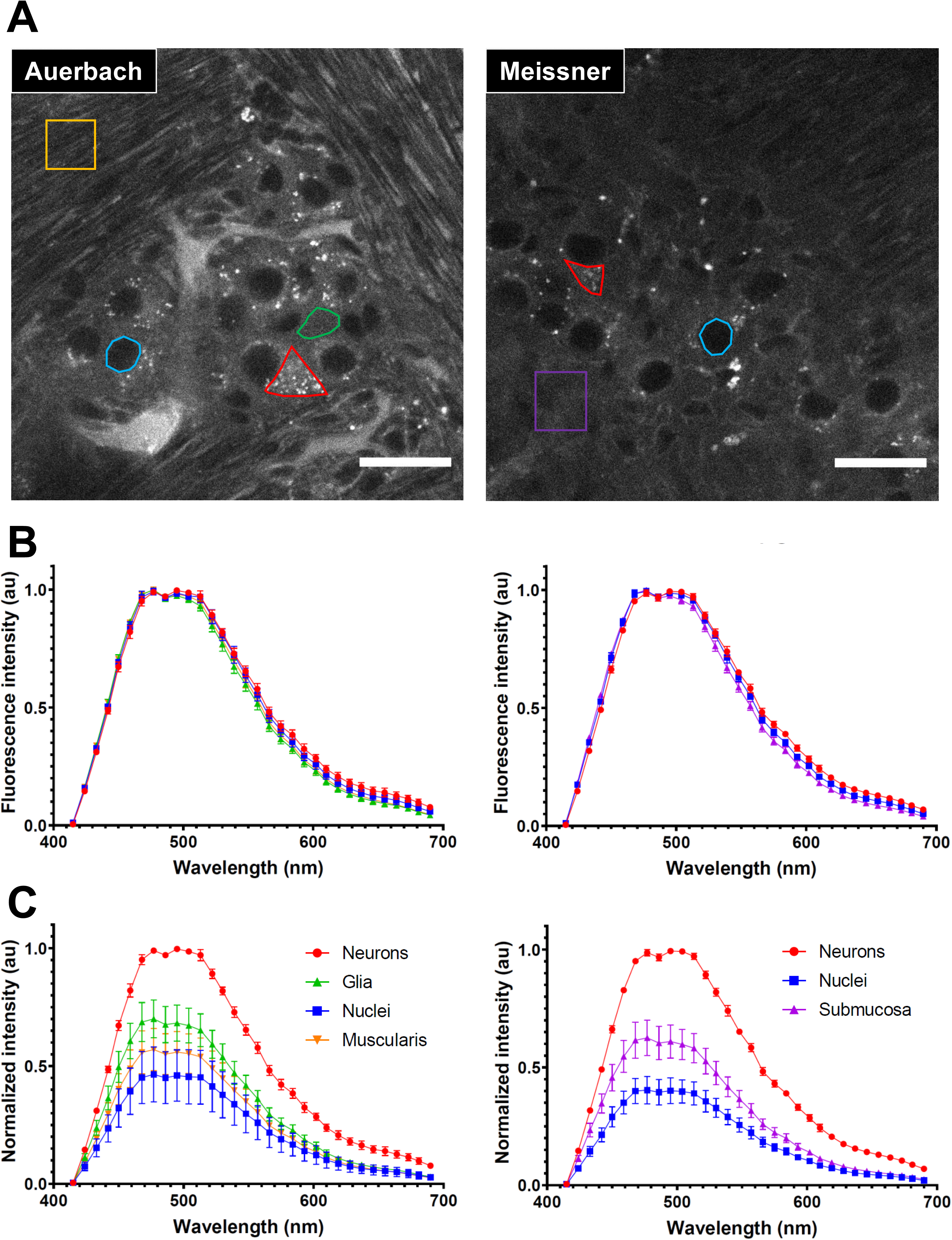
Characterization of the autofluorescence of the ENS. (**A**) Autofluorescence images taken at the level of the Auerbach’s (left) and Meissner’s (right) plexuses. Representative regions of interest used for spectral characterization are highlighted in color (red: neurons cell bodies, green: enteric glia cytoplasm, blue: neurons nuclei, yellow: muscularis, purple: submucosa). Scale-bars: 25 µm. (**B**) Spectral signature of the regions of interest depicted in A. at the level of the Auerbach’s (left) and Meissner’s (right) plexuses. (**C**) Relative fluorescence intensity of the regions of interest depicted in A. at the level of the Auerbach’s (left) and Meissner’s (right) plexuses. Error-bars: standard deviation.

At the level of Meissner’s plexus, ROIs were selected in 3 structures: neuronal cell bodies (red), neuronal nuclei (blue) and within the submucosa (purple) (**Figure 4A – right**). As previously observed in the Auerbach’s plexus, these regions had indistinct AF emission spectra (**Figure 4B – right)** and they showed similar differences in terms of AF intensity (**Figure 4C – right**).

Based on these observations, we conclude that the AF arising from the ENS does not exhibit a strong spectral variability, allowing its broad-band detection to maximize the collection efficiency. However, the strong intensity contrast observed between different structures of the ENS allows a relatively straightforward identification of cellular and subcellular structures.

### Two-photon spinning-disk microscopy allows fast, deep and resolved imaging of non-labelled ENS

3-D imaging on single-flying spot laser-scanning confocal microscopes is time-consuming and repeated scans to obtain z-stacks induced AF photobleaching (not shown). Two-photon excitation fluorescence (2PEF) microscopy strongly reduces out-of-focus photobleaching as excitation only occurs at the focal plane of the objective, but it cannot significantly reduce the acquisition duration. Recently, we introduced a new kind of spinning-disk 2PEF microscope based on a single spinning-disk and efficient light detection (Rakotoson et al., 2019). Similar to a 2PEF light-sheet microscope, our instrument produces a thin, planar non-linear excitation, and with the here used objective and detector, we could acquire a 150×150 µm AF image at a 182-nm pixel size in 480 ms. With a spatial resolution close to the one obtained on a confocal microscope (around 330 nm) and a temporal resolution 10 times faster, our OASIS microscope is particularly adapted to studying the intricate 3-D structure of the ENS.

We imaged the AF of an entire cleared intestinal wall, starting from the external face down to the crypts (**Figure 5** and **Supplementary Video S1**). Both muscularis longitudinalis (**Figure 5A**) and circularis (**Figure 5C**) could be visualized and recognized by their perpendicular organization of the muscle fibers. In between, Auerbach’s plexus (**Figure 5B**) exhibited a strong AF contrast and ganglia could be readily identified. Neurons displayed intense AF granules in their cell bodies (**Figure 5B - arrow**) in comparison to surrounding enteric glia (**Figure 5B - arrowhead**). Meissner’s plexuses were characterized by ganglia that were composed of systematically fewer neurons compared to Auerbach’s plexuses (**Figure 5D - arrow**). As a consequence of the non-linear excitation at 800 nm, a Second-harmonic Generation (SHG) signal can also be detected in the submucosa, arising from fibrillary structures. Crypts appeared as contrasted structures with sparse AF granules that have earlier be identified as zymogen-rich granules in Paneth cells (**Figure 5E**). An orthogonal (xz) reconstruction of the intestinal wall (virtual transversal slice) clearly identifies the different layers of the intestinal wall throughout its entire volume (**Fig. 5F**). Thus, the combination of rapid tissue clearing and fast 3-D AF imaging on the OASIS microscope is of particular interest for the morphological study of a sparse and inhomogeneous structure like the ENS and the label-free imaging allows for a surprisingly detailed analysis of the different layers and cell-types of the enteric system.

**Figure 5:**
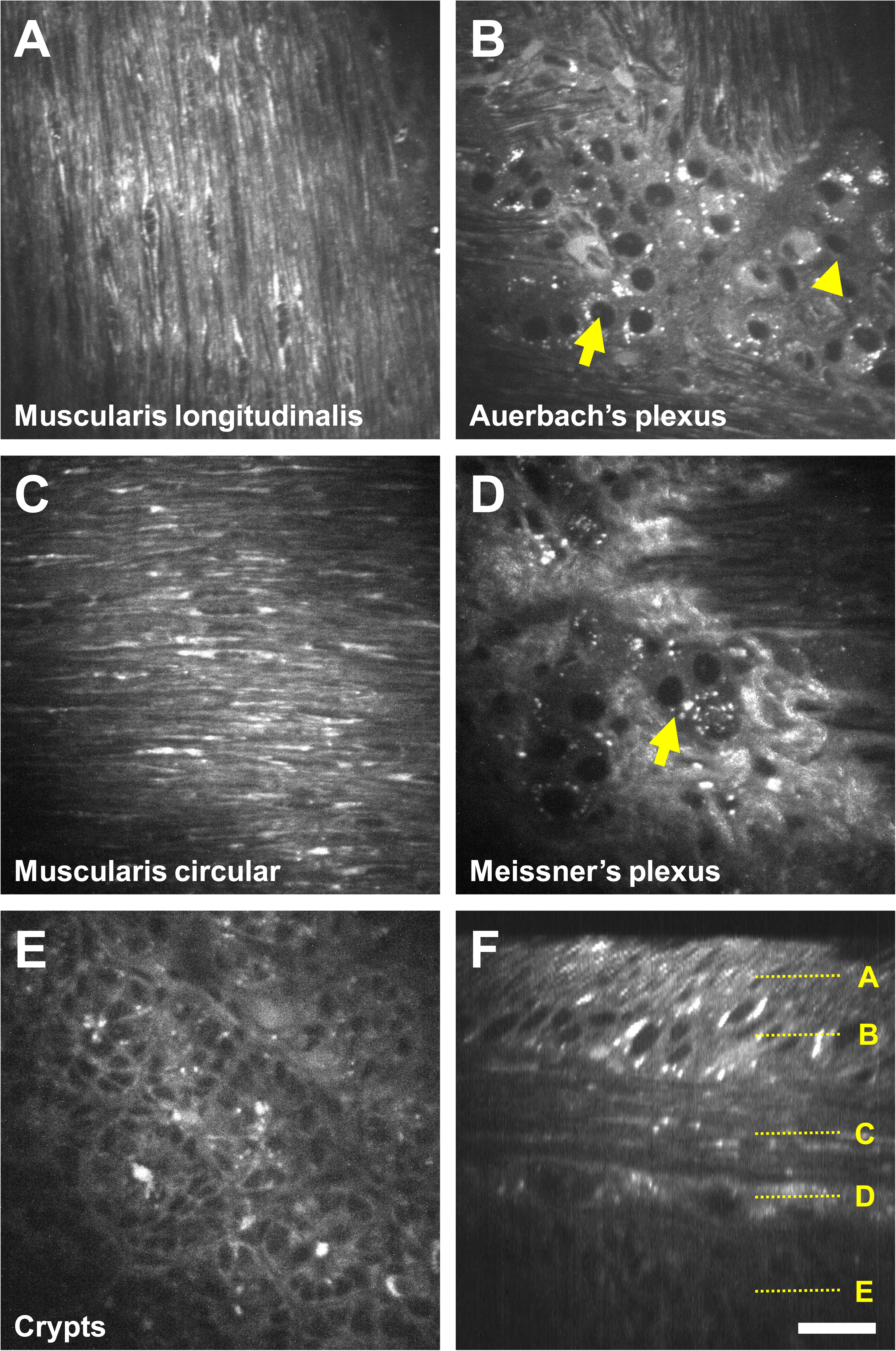
Fast 3D imaging of the ENS on the OASIS microscope. (**A-E**) Autofluorescence images taken under an 800 nm excitation at different depths of the intestinal wall starting from the muscularis longitudinalis to the crypts. Arrows: neurons, arrowheads: enteric glia. (**F**) Orthogonal reconstruction based on a z-stack acquired with 1 µm steps. Localizations of images A to E are depicted on the right. Scale-bar: 25 µm.

## DISCUSSION

In this study, we combined an innovative clearing method that can be geared to preserve different autofluorescent components with the unique performance of our OASIS 2P-microscope to demonstrate fast volumetric label-free imaging of the ENS. Our main findings are, (1), label-free imaging permits the unequivocal identification of both Auerbach’s and Meissner’s plexuses, validated through immunofluorescence controls experiments on small intestine sections; (2), by charactering ENS AF properties (fluorescence emission spectra upon 405 nm (1P) or 800 nm (2P) excitation) we unambiguously identified neuronal and glial cell types; (3), we applied fast clearing protocols on small intestine rings and reconstructed nervous plexuses in 3-D at high spatial resolution (300 nm laterally, 1.5 µm axially) and high throughput (2 images/s) on the OASIS virtual light-sheet microscope. Our study underpins the robustness of intrinsic biological signals for a detailed 3-D imaging of the ENS.

As of today, ENS is mainly studied using conventional histological slices that are stained or immunolabeled. As already discussed in the introduction, such approach is not optimal for studying a mesh-like and intricate structure like the ENS. Different approaches were proposed to overcome this problem. Among them, the whole-mount technique (Lebouvier et al., 2010a) enables a careful dissection of the intestinal wall to isolate parts of the ENS that can be immunolabeled and observed. This approach allows the observations of a few ganglia and their connections but it remains highly time-consuming, requires a strong expertise and it is not free of artifacts. Furthermore, whole-mount preparations isolate the ENS from other structures, the visualization of the interactions and spatial relationships in between different components of the intenstinal wall is thus impossible. Moreover, the dissection of the ENS, a tedious and highly delicate process, can lead to tissue damage. Finally, once extracted from the surrounding tissue that support its structure, the ENS 3-D architecture is modified. Thus, such protocol is not optimal for morphological studies. Instead, *in vivo* experiments were proposed, such approaches require the use of transgenic animals that undergo complex surgery and imaging protocols (Rakhilin et al., 2016). This can be particularly suitable for physiological experiments (Boesmans et al., 2019) but not for studying the 3-D morphology of the ENS at large scale and with a subcellular spatial resolution in a short amount of time. Based on these findings, the development of a complete workflow allowing to study the 3-D architecture of the ENS in its environment is of particular interest. That is why we have proposed the methodology described in the present paper that is based on, (1) a fast, harmless and easy clearing method to render the whole intestinal wall transparent, (2) a label-free approach based on tissue AF and (3) a new innovative microscope.

Tissue samples are not transparent. Scattering limits the penetration of light that is deviated from its original trajectory with a mean-free path <100 µm between two scattering events. To overcome this limitation and increase the maximal imaging depth in optical microscopy, clearing methods have been developed (Richardson and Lichtman, 2015; Azaripour et al., 2016; Seo et al., 2016; Ariel, 2017; Yu et al., 2018). The aim of such approaches is to make tissues more transparent by removing scattering centers and matching the refractive index (RI) of its components and hence improving both excitation penetration and fluorescence imaging. Various approaches were proposed and successfully tested: (1) organic solvents that extract lipids and homogenize the RI to a high value (fast technique but use of hazardous chemicals ; *e*.*g*. BABB, uDISCO), (2) high RI aqueous solutions that match the average RI (simple, non-hazardous but sometimes slow ; *e*.*g*. TDE, ClearT2, RapiClear^©^), (3) hyperhydrating solutions (simple, non-hazardous but extremely long ; *e*.*g*. Scale, CUBIC) and (4) hydrogel embedding (excellent clearing but slow and hazardous ; *e*.*g*. CLARITY). In this context, we have developed a new ultra-rapid, harmless and efficient clearing method based on a high-RI aqueous solution. Our approach, named BRG4 cannot be described here, because of the ongoing patent process. However, we have characterized and highlighted its advantages for clearing whole intestinal samples. Notably, we demonstrated that the plateau of transparency (around 75%, quantified in terms of visibility of a stripe pattern) was reached in less than 15 min. The level of transparency reached, comparable or higher to what is attained by other techniques means that the intestinal sample is not fully, but enough, transparent to perform microscopy across the entire intestinal wall (from the serosa to the crypts) even on a conventional laser-scanning confocal microscope with the pinhole closed to 1 Airy Unit (AU). Other clearing approaches can generate a higher transparency, however they require much more time and they are often hazardous for the operator or corrosive to the microscope objective, excluding the use of dipping lenses. Also many clearing protocols include a depigmentation step before the RI matching. In the case of a mildly pigmented organ like the small intestine we demonstrated that this step is dispensable. No differences in term of transparency after a 15 min immersion in the clearing solution was observed in depigmented samples. Only a small difference was observed for the half-life (less than 3 min), which is negligible in the light of the 45 min-long depigmentation step. Moreover, we have also observed a decrease of the AF signal after depigmentation. Thus, avoiding depigmentation, (1), saves time in the protocol and, (2), preserves AF intensity.

AF arises from various biomolecules (elastin, lipofuscin, …) (Zipfel et al., 2003) and is often considered as a source of background signal in immunolabeling experiments, that is why protocols and various reagents were developed for AF removal. However, various studies have highlighted the potential of this intrinsic signal for obtaining contrast and perform label-free microscopy in various organs and tissues including the gut (Ricard et al., 2012). Despite a low quantum yield, AF has often a large emission spectra enabling a collection over a wide range, thus increasing the signal collected. AF of the gut was reported upon mono-(Bhattacharjee et al., 2018) and two-photon excitation (Ricard et al., 2012) in healthy conditions but also for studying pathologies like Hirschsprung’s disease (Aggarwal et al., 2013). Here we used AF to image the ENS and demonstrated that the contrast obtained in our images is not based on a difference of AF emission spectra but rather on a difference of AF intensity of the various structures of the intestinal wall. Our investigations show that ENS components including Auerbach’s and Meissner’s ganglia, connections in between them, neurons and enteric glia can reliably be discriminated based only on AF. We confirmed this specificity by immunolabeling against HuC/D, a pan-neuronal marker commonly used to study the ENS (Mazzoni et al., 2021), and against S100β, a more reliable marker for enteric glia than the often used GFAP(Grundmann et al., 2019). Interestingly, the intensities of AF and immunolabelling are commensurable so that both signals can be acquired on the same sample. In this case, AF signal can be used as a “counterstain” that gives morphological context information and immunolabelling highlights specific structures of interest. This dual approach increases the information content extracted from a fluorescence image. This can be of particular interest to study the precise localization of tiny structures in the complex 3D environment of the intestinal wall. A “counterstain” approach is commonly used in pathology labs using conventional histological stains and an immunolabeling protocol revealed by an enzymatic color reaction. This conventional approach can only be used on tissue slices (2-D structures) but not on intact tissues or organs (3-D structures). The ombination of both AF and immunolabelling lets itself in a natural way to full, 3-D acquisitions. Immunolabelling is also compatible with our clearing protocol enabling observations of labeled structures also in thick samples. This was demonstrated by the HuC/D immunolabeling of neurons of both Auerbach’s and Meissner plexuses located at different depths in the intestinal wall. AF can be obtained under mono-and two-photon excitation. Using the latter approach, it was previously demonstrated that SHG imaging can also be realized on intestinal sample (Ricard et al., 2012). SHG that only occurs under a two-photon excitation is a non-linear scattering process and enables the acquisition of a signal arising from certain non -centrosymmetric molecules such as Type-I and -III collagen that can be found in connective tissue (Friedl et al., 2007). SHG signal arising from the submucosa was observed during our acquisitions (see Fig 5D) and can be of valuable interest for morphological observations (*e*.*g*. to highlight fibrosis in pathological samples (Ranjit et al., 2016)). Also, it can be separated from AF due to its relatively narrow spectrum at half the fundamental wavelength. In contrast, AF detection is facilitated by the large emission spectra. It does not require specific adaptations nor adding costly instruments on the optical setup and can be performed on any fluorescence microscope with a sufficiently sensitive detector.

We demonstrated our approach on different microscopes including a conventional laser-scanning confocal microscope and on a prototype of a single-spinning disk two-photon microscope (OASIS). Laser-scanning confocal microscopes are widely available on imaging platforms but they are not designed for deep-tissue imaging. Repeated laser-scans at various depth in the sample are time consuming and, probably worse, they induce photobleaching. Despite its limitations, we demonstrated that confocal microscopy can be used for AF 3-D imaging of the ENS but at the cost of lengthy acquisition times (more than 6 s for each AF image plane in Fig 3). That kind of microscopy thus suitable for occasional 3D acquisitions is not recommended for imaging important volumes, large cohorts or for the fast 3-D examination of intestinal samples in pathology labs.

Laser-scanning two-photon microscopy was also proposed for morphological study of the intestinal wall. Contrasted images were obtained with the identification of the various layers and cell types (Ricard et al., 2012). This approach is more suitable for deep-tissue imaging as out-of-focus photobleaching is strongly reduced and signal collection is more efficient thanks to the absence of a pinhole and owing to the properties of two-photon excitation (Oheim et al., 2001). However, it still requires long scan times (4,5 s/image)(Ricard et al., 2012). Recently, we have developed a new large-field excitation two-photon microscope based on a single spinning-disk allowing considerably faster acquisition times (480 ms/image). This OASIS microscope was successfully tested for 3D imaging of brain organoids (Rakotoson et al., 2019) and its advantages for 3D AF imaging of the ENS were confirmed in the present paper. We have imaged the whole intestinal wall (140 µm with a 1 µm z-step, from the serosa to the crypts) in less than 1 min 30 s. Such performance is fully compatible for large scale study of the ENS on important volumes or in numerous samples. One would argue that such characteristics can also be obtained on light-sheet microscopes. However, despite close performances in term of temporal resolution, spatial resolution is lower in comparison to the OASIS. Moreover, the geometry of the optical setup in light-sheet microscopes requires a specific positioning of the sample for the orthogonal illumination that can be tedious, this is not the case for the OASIS as the excitation and emission pathways are the same.

Using the combination of these 3 approaches (clearing, AF and single spinning-disk two-photon microscopy), we have demonstrated a fast, 3D morphological imaging of the ENS that cannot be obtained with other conventional approaches. Contrast obtained with the AF signal allows a specific discrimination of various structures including the identification of neuronal bodies with dense granule-like structures. Morphological differences in between the Auerbach’s and Meissner’s plexuses can be highlighted and the various connections in between the ganglia can be tracked. Such level of details and 3D organization cannot be obtained with previously described methods. Moreover, artifacts due to the slicing process or dissection of the intestinal wall during whole-mount preparation are strongly reduced with our approach. Applications are numerous. First, for fundamental research studies on the organization of the ENS and its relationship with its environment and with the central nervous system. That kind of description can only be obtained using high-throughput 3D imaging methods on a consequent number of samples. Second, in the field of pathology to better understand the link in between neurodegenerative diseases such as Parkinson’s disease and the ENS (Lebouvier et al., 2010b; Ohlsson and Englund, 2019).

## MATERIALS AND METHODS

### Ethics statement

All experiments were performed in accordance with the French legislation and in compliance with the European Community Council Directive of November 24, 1986 (86/609/EEC) for the care and use of laboratory animals. The used protocols were approved by the local ethics committee. In total, this study used 12 C57BL/6 mice (Charles River) that were bred in the local animal house on an inverted dark-light cycle with access to food and water *ad libitum*.

### Sample preparation

#### Intestine extraction and fixation

Mice were euthanized with carbon dioxyde and their small intestine was dissected out. Intestines were then flushed and washed with a solution of 1X Phosphate-Buffered Saline (1X PBS) at pH 7.2 (Gibco) and then divided into 3 parts: *duodenum, jejunum* and *ileum* and fixed overnight (ON) at 4°C in formalin (VWR), a 4% formaldehyde solution buffered to pH 6.9. The thus cleaned samples were stored until use in 1X PBS/0.02% NaN_3_ at 4°C. All of the experiments were realized on ileum sections.

#### Freezing and cryostat section of samples

Fixed tissue was cut into 1 cm fragments and first impregnated during 3h in a 15%-sucrose solution (Merck) in 1X PBS, 0.02% NaN_3_ and placed thereafter in a similar solution, but containing 30%-sucrose solution for cryoprotection. Samples were embedded with Optimal Cutting Temperature (OCT) (CellPath) and immersed in ice-cold isopentane in dry ice. Once frozen, 7-µm thin transversal slices were cut on a cryostat (Reicher-Jung, Cryocut 1800; Leica) and deposited on positively charged BK-7 glass slides (Menzel-Gläser Superfrost Plus; Gerhard Menzel). Slides were conserved at -80°C until use.

#### Swiss Roll

Fixed mouse intestinal tissues were incised longitudinally to expose the lumen (Bialkowska et al., 2016). The sample was processed as described in the previous section and – after the cryoprotective step – using a wooden toothpick, the samples were rolled up on themselves, the luminal side facing the inside and placed in plastic moulds filled with OCT and frozen at - 80°C. Finally, 7-µm thick cryostat sections were cut.

### Staining, immunofluorescent labelling and clearing

Nota: Due to the long incubation times in each protocol, all reagents used contain 0.02% NaN_3_ as preservative.

Room temperature (RT) in defined as between 20°C and 25°C (France).

#### Hematoxylin/Erythrosin/Saffron (HES) staining

Once extracted, intestines were washed in PBS and fixed overnight in a 4% formaldehyde solution (Merck). Then, they were stored in ethanol 70% before the embedding process. Tissues are dehydrated in increased concentration of ethanol, then isopropanol and finally impregnated in paraffin (Logos One, Thermo Fischer), before embedding. Slides were deparaffinized and stained with different morphological reagents for HES (Hematoxylin/Erythrosin/Saffron): nuclei were stained with hematoxylin in blue, then cytoplasms in pink with erythrosin and finally collagen fibers in yellow with saffron. Staining was realized in Spectra, Leica automaton. Slides were stained and dehydrated in automaton (SPECTRA, Leica), then mounted in a permanent mounting medium (Pertex).

#### Thick-sample Immunolabeling

The previously fixed tissues were cut into 1.5-cm long sections. An antigen retrieval step by successive methanol baths ensuring a dehydration / delipidation of the cells was performed at RT, followed by a permeabilization step with an aqueous solution of 0.2% Triton X100 (Bio-Rad Laboratories), 0.2M Tris pH8, 20% DMSO and 0.3M Glycine for 1h followed by a blocking step using an aqueous solution of 0.2% Triton X100 (Bio-Rad Laboratories), 0.2M Tris pH8, 10% DMSO and 10% goat serum for 5h, all under agitation and at 37°C. Tissue slices were incubated with a primary antibody anti-HuC/D (1/250; Ab184267; Abcam) or anti-S100β (1/100; ab52642; Abcam) in an aqueous solution of 10% washing buffer 10X, 10% goat serum and 5% DMSO in Eppendorf tube for 40 h under agitation at 37°C. After 5 successive washes of 30 min and 1 wash of 1 h under agitation at RT with an aqueous solution of 0.2M Tris pH8 and 0.2% Tween 20, the samples were incubated with a secondary antibody (Goat anti rabbit conjugated Alexa fluor 633; 1/1000; A-21070; Invitrogen) in an aqueous solution of 10% washing buffer 10X, 10% goat serum and 5% DMSO for 20 h, under agitation, in the dark at 37°C. The samples were then washed as before were and protected from light.

#### Thick-sample clearing

Clearing is the result of delipidation and RI (refractive index) homogenization (ref). The exact formulation cannot be released at this time, DPG4 and BRG4 and process are under patent deposit.

DPG4, a depigmentation solution, contain hydrogen peroxide.

BRG4 contain a mixture of detergent, buffer, and 60% TDE (2,2’-thiodiethanol). The refractive index of the mixture is about 1.47. Clarification of the small intestine is achieved in 20 min at 37°C. Clearing is achieved in one step.

### Imaging

#### Brightfield

Images of the HES staining were taken on a Axioscan Z1 (Zeiss) with a 40x objective.

#### Macroscope

Images illustrating the clearing steps of thick intestinal sample were obtained with the SMZ800 stereo macroscope, equipped with a digital camera DS-Fi and DS-U2/L2 USB (Nikon). Samples were illuminated with white light (3000°K) and the images acquired with the NIS-Element software.

#### Confocal microscopy

Images of tissue sections and thick samples were acquired on a LSM880 inverted confocal and LSM710 upright laser scanning microscopes (Zeiss). Autofluorescence (AF) images were obtained under a 405 nm excitation wavelength and immunolabeling images under 633nm excitation. The objective used was a 40x/NA1.2 water-immersion objective (Zeiss Plan-Apochromat) and all images were acquired with the pinhole opened at 1 AU. For AF acquisitions, photons were collected in between 417 and 740 nm. For immunolabeling acquisitions signal was collected in between 415 and 599 nm and AF in between 645 and 735 nm.

#### Non-linear microscopy

The OASIS microscope is described in detail in (Rakotoson et al., 2019). Briefly, 2P images were acquired upon 800 nm two-photon excitation and the autofluorescence signal was collected in between 400 nm and 568 nm. Acquisition time was set to 480 ms for each image.

### Image analysis, quantification and statistics

#### Image processing

Images were visualized and analyzed with the ImageJ/Fiji software. Data were processed with the Excel (Microsoft) and Prism (GraphPad) softwares.

#### Clearing analysis

1.5 cm-long intestinal sections were separated in two groups. The first group (n=6) was depigmented during 45 min with an in-house DPG4 solution (patent pending); the second group (n=4) was used as a control and was not depigmented. Samples were placed in a home-made tank with a black and white pattern located below its glass bottom. Images were acquired on a SMZ800 stereo macroscope (Nikon) as described above. A first image was acquired with the sample immersed in a 0.2M Tris solution as a reference. The Tris solution was then removed and replaced with custom BRG4 clearing solution (patent pending). After the immersion of the sample in the clearing solution, images were acquired at various timepoints (every 10 s during 5 min, every 30 s during 5 min and finally every minute during 10 min. For each acquisition, the Michelson contrast (REF) of the intestinal sample on the black and white pattern was measured at each timepoint. Data were then processed on the Prism (GraphPad) software and the half-life and plateau values were extracted for each conditions (depigmented *vs*. non depigmented).

#### Spectral analysis

Autofluorescence spectra were acquired from non-depigmented cleared intestinal samples. Spectral images were taken at the level of the Auerbach’s and Meissner’s plexuses using the 32-PMTs spectral PMT array of a LSM880 confocal microscope (Zeiss) upon 405nm excitation. At the level of the Auerbach’s plexus, regions of interest (ROI) were selected in the cytoplasm and nuclei of neurons, in the cytoplasm of enteric glia and in the muscularis. At the level of Meissner’s plexus, ROI were selected in the cytoplasm and nuclei of neurons and in the submucosa. Mean fluorescence intensity was extracted in each ROI and for each of the 32 PMTs before normalization using the ImageJ software. Data are from 8 and 6 independent acquisitions at the level of Auerbach’s and Meissner’s plexuses, respectively.

### Statistics

All other results are at least triplicates of three independent experiments and are represented as mean ± SD. Mann-Whitney test was used to compare among experiments.

## Acknowledgements

The authors thank the HistIM platform “Histologie, Immunomarquage, Microdissection laser” of Institut Cochin (INSERM U1016, CNRS UMR 8104, Université Paris Cité) for their precious help for the HE experiments. Fabrice Licata (Imaging platform) and Thierry Bastien (Prototyping platform) of the UMS BioMedTech facilities (Université Paris Cité, CNRS UMS2009, INSERM US36) are also acknowledged for their contributions.

